# Context-specific interaction of the lipid regulator DIP-2 with phospholipid synthesis in axon regeneration and maintenance

**DOI:** 10.1101/2025.02.06.636954

**Authors:** Seungmee Park, Yishi Jin, Andrew D. Chisholm

**Affiliations:** Department of Neurobiology, School of Biological Sciences, University of California, San Diego, La Jolla, CA 92093, USA

**Keywords:** phosphatidylethanolamine, phosphatidylcholine, Kennedy pathway, aging, axon regeneration, axon degeneration

## Abstract

Neurons maintain their morphology over prolonged periods of adult life with limited regeneration after injury. *C. elegans* DIP-2 is a conserved regulator of lipid metabolism that affects axon maintenance and regeneration after injury. Here, we investigated genetic interactions of *dip-2* with mutants in genes involved in lipid biosynthesis and identified roles of phospholipids in axon regrowth and maintenance. CEPT-2 and EPT-1 are enzymes catalyzing the final steps in the *de novo* phospholipid synthesis (Kennedy) pathway. Loss of function mutants of *cept-2* or *ept-1* show reduced axon regrowth and failure to maintain axon morphology. We demonstrate that CEPT-2 is cell-autonomously required to prevent age-related axonal defects. Interestingly, loss of function in *dip-2* led to suppression of the axon regrowth phenotype observed in either *cept-2* or *ept-2* mutants, suggesting that DIP-2 acts to counterbalance phospholipid synthesis. Our findings reveal the genetic regulation of lipid metabolism to be critical for axon maintenance under injury and during aging.

**Article Summary:** Little is known about how adult neurons live long with limited regenerative capacity. This study investigates the role of lipid metabolism in sustaining neuronal health in *C. elegans.* Mutating phospholipid synthetic genes impairs axon regrowth after injury. Lack of DIP-2, a lipid regulator, restores regrowth, suggesting DIP-2 counterbalances phospholipid synthesis. Moreover, neuronal phospholipid synthesis is essential for preventing age-dependent axonal defects. These findings reveal phospholipid biosynthesis is key to axon integrity during aging and injury. As lipid metabolism is implicated in neurological disorders, this study serves as an entry point into investigating neuronal lipid biology under various conditions.

## Introduction

Lipid metabolism has gained growing attention for its implications in neurological disorders (Tracey *et al*. 2018; Rickman *et al*. 2020). In the nervous system, lipids are critical as membrane building materials, the primary component of myelin, lipid-based neurotransmitters, regulators of synaptic plasticity, and signaling molecules (Bazan 2003; Cermenati *et al*. 2015). Phospholipids are one of the major constituents of cellular membranes and therefore their biogenesis and asymmetric distribution across the membrane are crucial for neuronal integrity (Vance 2015). The elaborate morphology of adult neurons is maintained by both intrinsic and external factors. Aging causes a reduction in phospholipid levels in the brain (Hancock *et al*. 2022) and renders neurons increasingly vulnerable and less resilient in response to injury and stress (Mattson and Magnus 2006; Wang and Michaelis 2010; Popa-Wagner *et al*. 2011; Mcewen and Morrison 2013). However, our understanding of how these effects affect neurons remains limited.

In eukaryotic cells, phosphatidylcholine (PC) and phosphatidylethanolamine (PE) are the two major types of phospholipids found in cellular membranes (Suetsugu *et al*. 2014). They can be up-taken from the environment via ingestion of food and also formed by several biosynthetic pathways (Watts and Ristow 2017). The Kennedy pathway synthesizes both PC and PE *de novo* through three sequential enzymatic steps in two parallel branches (Gibellini and Smith 2010). Deletion of pcyt1a, one of the two genes encoding the rate-limiting factor CCT in the PC branch, leads to early embryonic lethality in mice (Wang *et al*. 2005), demonstrating that the *de novo* biogenesis is required for early development. Studies have shown that phospholipids can be synthesized locally in axons of cultured mammalian neurons (van der Veen *et al*. 2017). In *C. elegans*, deficiency of the Kennedy pathway enzyme *cept-2* results in reduced axon regrowth following injury (Kim *et al*. 2018). In mice, tissue-specific conditional loss of function in Kennedy pathway genes has little effect on optic nerve regrowth in a wild type background but reduces regrowth in backgrounds such as *Dgat1* KO or *PTEN* KO (Yang *et al*. 2020a). Conversely, overexpression of PCYT genes in a tissue-specific manner can enhance axon regrowth (Yang *et al*. 2020a), consistent with PC or PE being rate-limiting in axon regrowth across species.

*C. elegans* is a tractable model suited for studying the effect of altered phospholipid biosynthesis on neurons under injury and aging conditions *in vivo* due to short lifecycle and lifespan. It encodes orthologs of all enzymes in the Kennedy pathway (Watts and Ristow 2017). Inhibition of the Kennedy pathway enzymes PCYT-1 or CEPT-1 results in reduced PC levels but normal PE (Walker *et al*. 2011), indicating that indeed phospholipids are synthesized in *C. elegans*. In addition to the *de novo* steps, both PC and PE can be synthesized through the salvage pathways: PC from phosphoethanolamine by methyltransferase enzymes PMT-1 and PMT-2 and PE from phosphatidylserine (PS) in mitochondria by PSD-1 (Wang *et al*. 2014). Loss of function in PSD-1 results in lethality, consistent with findings from other organisms that the Kennedy pathway and PSD pathways are not redundant at the cellular or organismal level (Steenbergen *et al*. 2005). Studies have revealed that the Kennedy pathway is required for various cellular processes, including maintaining membrane fluidity at low temperatures to support cold adaptation in animals (Svensk *et al*. 2013) and preventing germline ferroptosis (Kim *et al*. 2018). It remains to investigate how this pathway is critical in neurons in response to injury and with age.

Here we report that *C. elegans cept-2* and *ept-1*, encoding the enzymes catalyzing the last step the Kennedy pathway, are independently required for axon regrowth and integrity. CEPT-2 acts cell-autonomously to maintain axon morphology especially with age, implying the critical role of phospholipid synthesis within aging neurons. Notably, lack of the lipid regulator DIP-2 fully suppressed the reduced axon regrowth defects of *cept-2* or *ept-1* null mutants. Our studies demonstrate the role of key lipid biosynthetic genes in neuronal resilience in response to injury and aging, and reveal DIP-2 as acting antagonistically to lipid biosynthesis.

## Results

### Choline/ethanolamine phosphotransferase CEPT-2 is essential for development and reproduction

Genetic redundancy exists in many steps of the Kennedy pathway (Figure 1A); the two paralogs *cept-1* and *cept-2* encode choline/ethanolamine phosphotransferases, which catalyze the last step of both branches where two fatty acids from diacylglycerol are transferred to choline/ethanolamine. Loss-of-function mutants of *cept-1(et10)* do not exhibit overt phenotypes (Svensk *et al*. 2013). The *cept-2(ok3135)* allele is a deletion predicted to cause frameshift and a premature stop codon (Figure 1B). >90% of these mutants are embryonically lethal and yet <5% are escapers that grow to adulthood (Figure S1A). We were unable to obtain double mutants of *cept-1(et10)* and *cept-2(ok3135),* suggesting double mutants may be inviable. We further generated a second allele *ju1669*, which deletes 4 nucleotides in the second exon and would result in a frameshift and a premature stop codon (Figure 1B). *ju1669* resembles *ok3135* in the fraction of viability (Figure S1A). These observations demonstrate that both *ok3135* and *ju1669* are null alleles; hereafter both will be indicated as *cept-2(0)*. Transgenic expression of a *cept-2* minigene rescued the developmental and reproductive defects (Figure S1A,B). These results reveal that *cept-2* is necessary for animal growth and propagation over generations.

**Figure 1.**
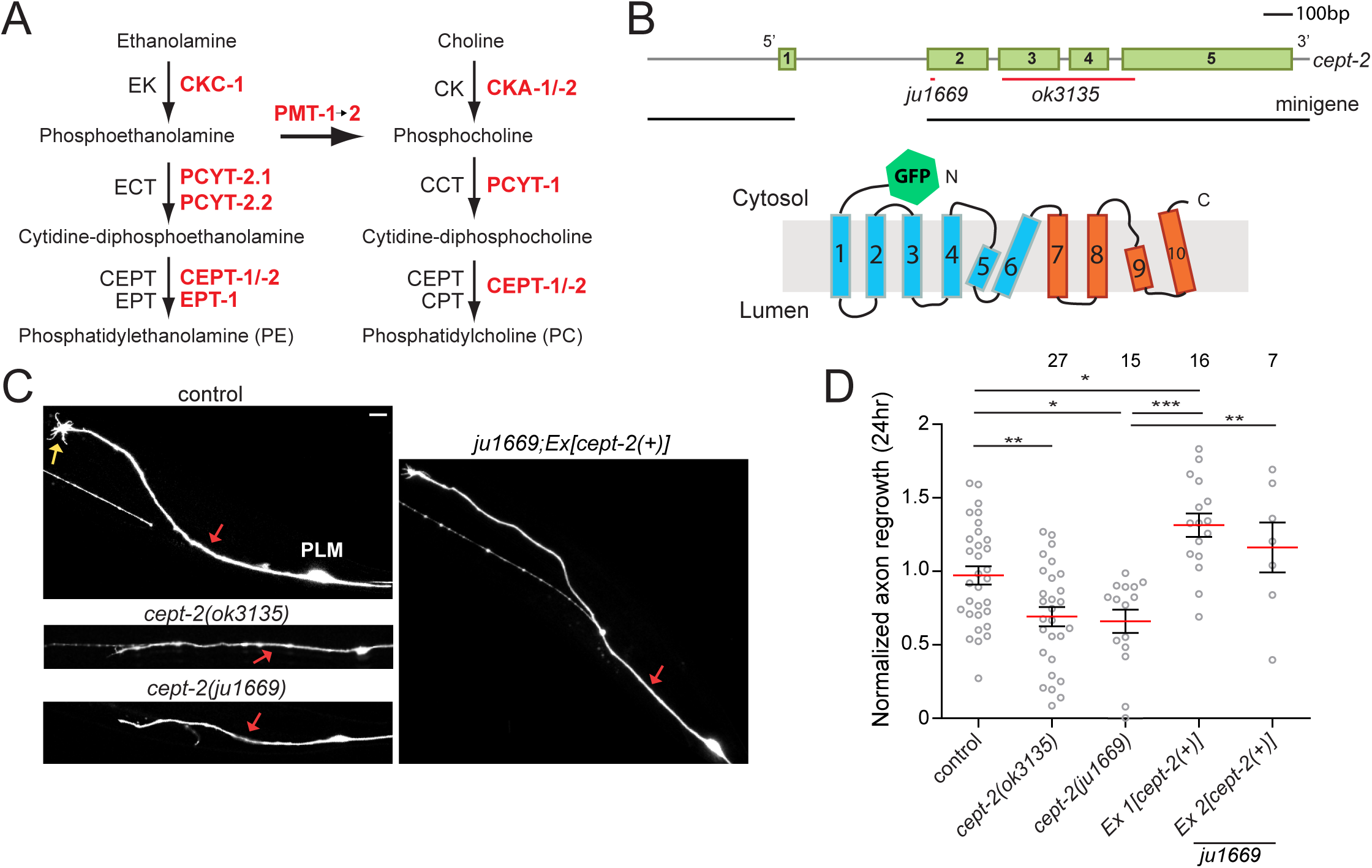
CEPT-2 is required for injury-induced axon regeneration. (A) The Kennedy pathway mediates *de novo* synthesis of phosphatidylethanolamine (PE) and phosphatidylcholine (PC) in two parallel branches of three consecutive, enzymatic reactions. The enzymes in black indicate mammalian orthologs whereas those in red are *C. elegans* orthologs, based on sequence homology. (B) Gene and protein structure of *cept-2*. *Top,* red underlines indicate lesions in the *cept-2* gene. Black underlines represent regions included in constructing the minigene to rescue the phenotype of *cept-2*. Bottom, predicted CEPT-2 protein, composed of 10 transmembrane regions based on the structure of the human CEPT1 (Choline/ethanolamine phosphotransferase 1)(Wang *et al*. 2023). The first 6 transmembrane regions (blue) function as the catalytic domain while the last 4 (scarlet) transmembrane regions are important for dimerization, based on structure of human CEPT1(Wang *et al*. 2023). (C) z-stacked confocal images of PLM neurons acquired 24 h following injury; *Pmec-4-GFP(zdIs5)* background. Yellow and red arrows indicate a growth cone and the point of where the axon was severed, respectively. (D) Axon regrowth at 24 h following axotomy, normalized to the same day control mean. Mann-Whitney U test for comparison between control and *ok3135* or *ju1669*. Kruskal-Wallis test and Dunn’s post hoc test for comparison of control, *ju1669*, and rescued groups. The number of PLMs tested for each genotype is indicated above the corresponding genotypes in the graph. Scale bar = 10 µm. \**p*<0.05, \*\**p*<0.01, and \*\*\**p*<0.001.

### CEPT-2 plays a role in axon extension following injury

Touch receptor neurons (TRNs), critical for sensing gentle touch, have been established as a model for investigating the mechanism of axon regeneration after laser axotomy (Yan *et al*. 2009; Chen *et al*. 2011; Kim *et al*. 2018). We previously found that *cept-2(ok3135)* mutants exhibited diminished axon regrowth 24 h post-axotomy in TRNs (Kim *et al*. 2018). We found *cept-2(ju1669*) mutants displayed equivalent axon regrowth defects that were fully rescued by *cept-2*(+) transgenes (Figure 1C,D). Reduced axon regrowth at the 24 time point could reflect defects in growth cone formation or in axon extension process (Chen *et al*. 2011). To distinguish between these two possibilities, we examined growth cone formation in PLM axon 24 h post-axotomy and found no difference in the percentage of growth cone formation between control and mutants (Figure S1C). This result suggests that *cept-2* is necessary for axon extension processes rather than growth cone formation.

### CEPT-2 cell-autonomously maintains axonal integrity in aging neurons

TRNs display simple and stereotyped morphology, allowing for tracking various age-associated morphological defects (Pan *et al*. 2011; Tank *et al*. 2011; Toth *et al*. 2012; Noblett *et al*. 2019). In *cept-2(0)*, axons subject to axotomy at the late larval (L4) stage did not exhibit any overt morphological defects compared to control (Figure 2A). Similarly, 1-day old mutant adults did not exhibit any severe abnormality in axon morphology compared to control. We then asked whether *cept-2(0)* displayed age-dependent morphological defects up to day 5 of adulthood (Figure 2A,B). Control (P*mec-4*-GFP) adults displayed blebs, defined as protrusions that appeared to be filled with a lower intensity of fluorescence, along the axon process and beaded or thin terminal ends, consistent with prior observations of age-dependent deterioration in neuronal morphology (Pan *et al*. 2011; Toth *et al*. 2012). In contrast, *cept-2(0)* mutant adults displayed more severe axonal morphology defects; the most striking one was round swellings or beadings with a diameter of ∼3 µm (Figure 2C), many of which were present in the distal axon (>100 µm anteriorly from the soma) and axon termini but seldomly observed in the proximal axon near the soma. Some beadings appeared to contain liquid without any fluorescence inside. We conducted longitudinal analysis to track the formation of these beadings from L4 to day 5 adult stages; a typical beading was visible at the day 2 adult stage, and grew in size with age (Figure 2D). While the progressive defects were found throughout the axon, the soma morphology appeared to be unaffected. Likewise, another pair of bilateral touch neurons ALMs in *cept-2(0)* exhibited beadings along the axon though these defects appeared at later stages (day 3 or 4) than in PLMs as PLMs are generally more prone to exhibit these beadings in *cept-2* mutants (Figure S1D). All the axonal defects in *cept-2* TRNs were significantly rescued by expression of the minigene (Figure 2A). These findings demonstrate that *cept-2* is required for maintaining axon integrity particularly in aging neurons. Single cell transcriptomic studies of L4 stage neurons suggest *cept-2* is expressed in both ALMs and PLMs (Taylor *et al*. 2021).

**Figure 2.**
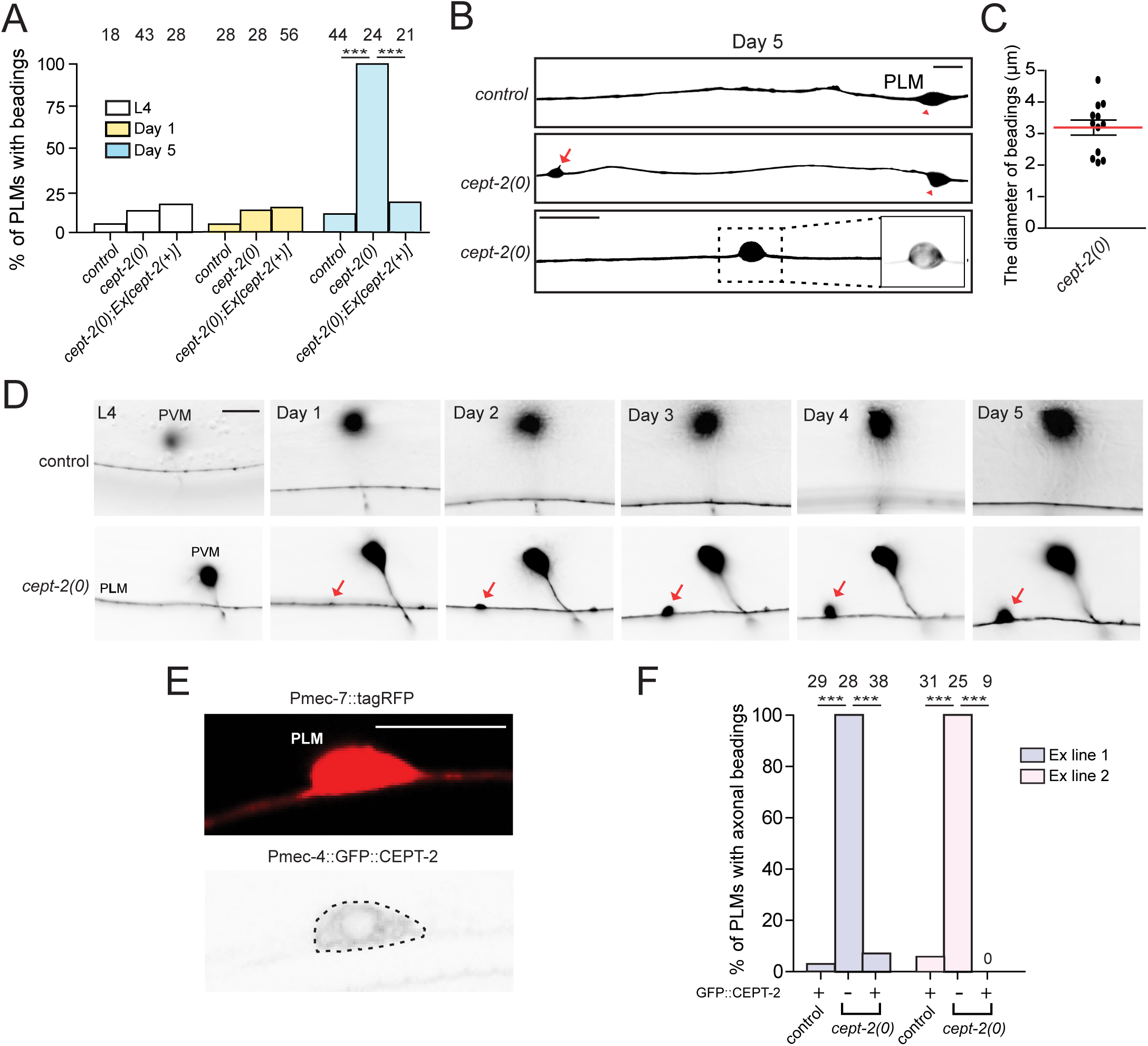
CEPT-2 maintains axonal integrity during aging. (A) Quantitation of axonal beadings with the average of ∼3 µm in diameter observed on PLM axons in the *zdIs5* background at L4 and 1- and 5-day old adult stages. The number of PLMs scored for each genotype is shown above the corresponding bar. Fisher’s exact test used for statistics. (B) z-stacked confocal images of PLM neurons (*zdIs5*) in 5-day old adults. Red triangles point to the PLM soma. Red arrow indicates a beading on the axon. The most bottom image shows the magnification of a beading with non-fluorescent interior. (C) Quantitation of the diameter of axonal beadings. (D) Longitudinal analysis of a beading formation on PLM axon in the *zdIs5* background. Red arrows point to the large beading imaged from the same animal from L4 to 5-day old adult stages. (E) *Top*, schematic diagram illustrates the PLM soma filled with red fluorescence. *Bottom*, single-plane confocal images of PLM labelled with *jsIs973 [Pmec-7-mRFP]* and expressing *juEx7992 [Pmec-4-GFP::cept-2 cDNA]*. (F) Quantification of PLMs with axonal beadings in 5-day old adults. *Pmec-4-GFP*::*cept-2* cDNA is expressed from ex line 1 (*juEx7993)* and 2 (*juEx7992)* in the *zdIs5* background. The number of PLMs scored for each genotype is indicated above the corresponding bar. Fisher’s exact test. \*\*\**p*<0.001.

To determine whether CEPT-2 functions cell-autonomously, we expressed GFP::CEPT-2 under the control of the TRN-specific *mec-4* promoter. In touch neurons, GFP::CEPT-2 formed tubular structures encircling the nucleus and displayed a network-pattern in the cytoplasm (Figure 2E), consistent with the known localization of CEPT proteins to the ER (Henneberry *et al*. 2002; Horibata and Sugimoto 2021). Transgenic expression of GFP::CEPT-2 rescued the axonal beading phenotype in *cept-2* mutants at 5-day old stage and its overexpression in control did not cause any adverse effects on touch neurons (Figure 2F). These results indicate CEPT-2 can function cell-autonomously to maintain axonal morphology.

### Ethanolamine phosphotransferase EPT-1 is required for development and reproduction

While CEPT-2 is common to both PC and PE branches, EPT-1 is the *C. elegans* ortholog of the ethanolamine phosphotransferase EPT1 (Figure 3A; Figure S2A), the enzyme specific to the final step of the PE branch (Figure 1A). We asked whether loss-of-function mutation of *ept-1* would have any effect on neurons. The *tm3093* deletion is predicted to cause a frame-shift and premature stop codon (Figure 3B, denoted *ept-1(0)*). *ept-1(0)* mutants are thin, short, and pale especially in the intestine, and produce significantly lower brood size compared to control (Figure 3C,D). Expression of an *ept-1* genomic construct containing a 1.4 kb region upstream of ATG (Figure 3B) completely rescued body size and fertility defects (Figure 3C,D). We further visualized expression of EPT-1 by tagging GFP at its N-terminal end (Figure 3B). Expression of this GFP::EPT-1 construct in *ept-1(0)* rescued brood size to the same degree or more effectively in brood size than the untagged version while causing no effect on body size (Figure 3C,D), suggesting that tagging the protein with GFP disrupts its function weakly. Together, these results demonstrate that *ept-1* is necessary for proper development and reproduction.

**Figure 3.**
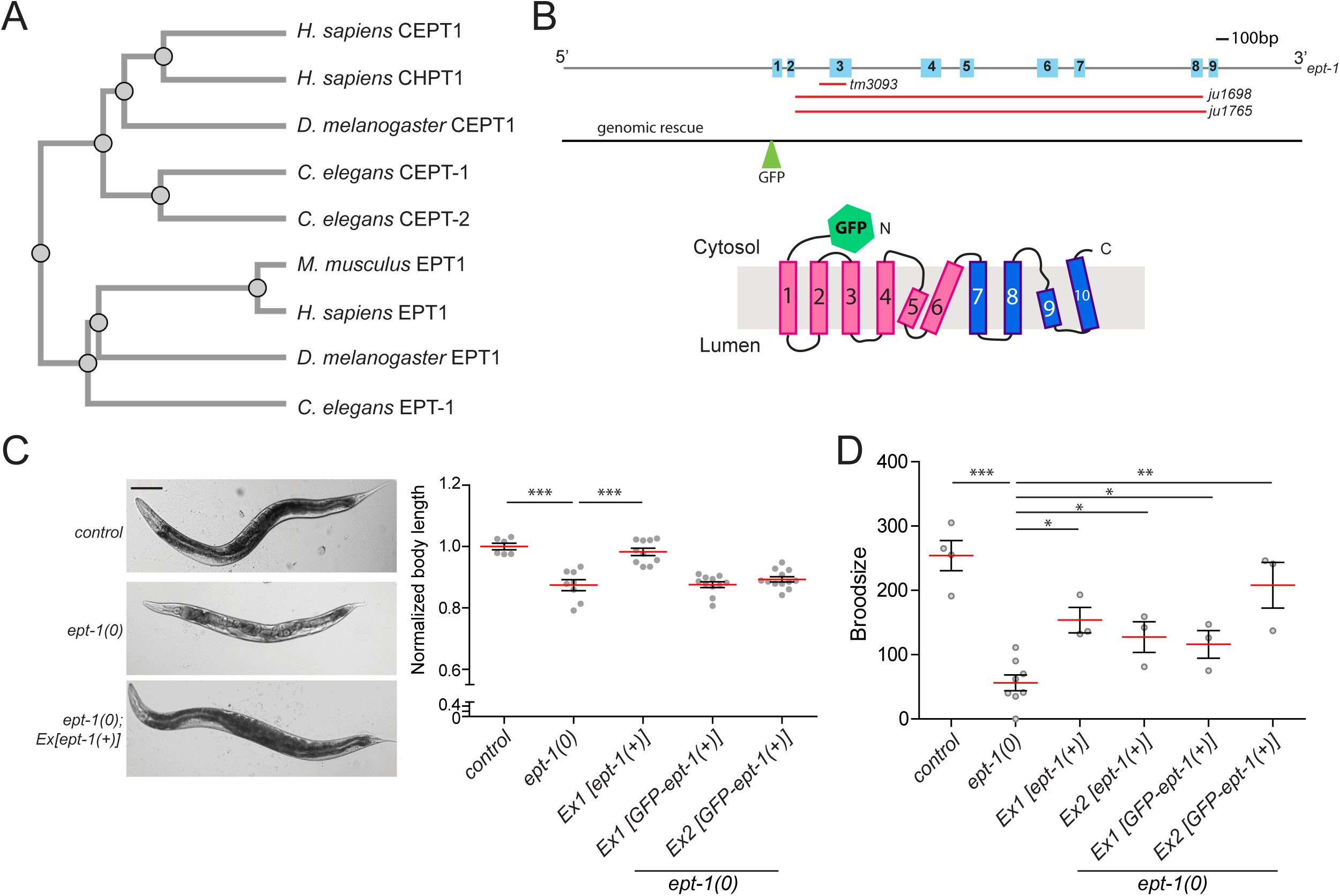
EPT-1 is required for development and reproduction. (A) Phylogenetic tree (Clustal Omega) illustrating divergence of EPT1 from CEPT1 and CPT1. (B) *Top,* gene structure of *ept-1*. Red lines indicate lesions in three different mutant alleles. Black line represents the region included in the genomic rescue construct *(juEx8002)*. *Below,* the predicted protein structure of EPT-1 tagged with GFP at N-terminus. (C) *Left*, brightfield images of control, *ept-1(tm3093)* mutant, and rescued *ept-1(tm3093)* mutant *(juEx8002)* worms. Scale bar = 100 µm. *Right*, quantitation of body length. 1-way ANOVA and Tukey’s post hoc test. (D) Mean brood size measured over 5 days of adulthood. Kruskal-Wallis test and Dunn’s post hoc test. \**p*<0.05 and \*\*\**p*<0.001.

### EPT-1 promotes injury-induced axon regrowth and maintenance of neuronal morphology in aged adults

*ept-1(0)* animals displayed largely normal PLM axon morphology at L4 stage (Figure S2B). We tested whether *ept-1* was required for PLM axon, and observed a reduction in axon regrowth 24 h post axotomy (Figure 4A); this phenotype was fully rescued by expression of the *ept-1* genomic construct (Figure 4A). Like CEPT-2, EPT-1 catalyzes the final step of *de novo* phospholipid synthesis, transferring fatty acids from diacylglycerol to ethanolamine. We therefore examined whether *ept-1(0)* mutants also displayed age-dependent defects in TRN morphology. *ept-1(0)* mutants displayed normal TRN morphology at the L4 and day 1 adult stages, except for the presence of very short (<1 µm) branches and blebs in ALM and PLM axons (Figure S2B). At day 5 of adulthood, we found a variety of morphological defects in ALM and PLM axons compared to controls, including ectopic branches, axon breakage or thinning, blebs, and irregularly shaped structures (Figure 4B; Figure S2C). The most striking phenotype was axon degeneration often observed in the middle and terminal regions of axons but absent in the proximal region near the TRN soma (Figure 4B). Overall, TRN soma shape was comparable to that of controls. Expression of either untagged or GFP-tagged genomic constructs significantly rescued axon degenerative defects in *ept-1* mutants (Figure 4B). We next asked where EPT-1 plays a role in maintaining axonal morphology. We found that GFP::EPT-1 expressed from a genomic rescue construct was strongly detected in the intestine, the spermatheca, and some cells in the head but not in the nervous system (Figure S2D). According to the single-cell RNA seq dataset from CeNGEN (Taylor *et al*. 2021), *ept-1* expression in the L4 stage is low in ALMs and undetectable in PLMs. EPT-1 may be expressed at very low levels or it may act non-cell autonomously to support the integrity of axonal morphology. Nevertheless, our results show that EPT-1 is critical for maintaining intact axon structures.

**Figure 4.**
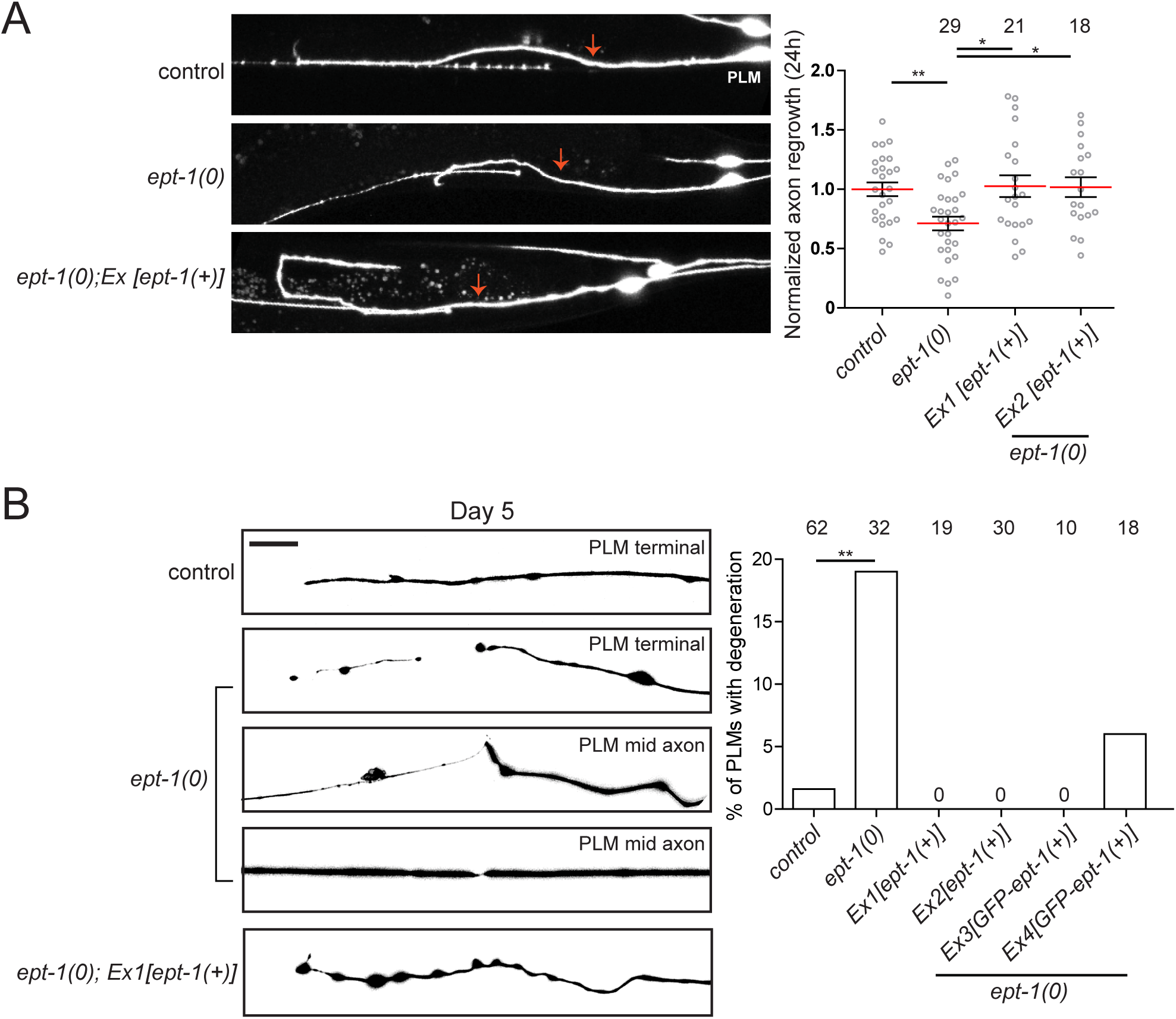
EPT-1 is required for both axon regeneration and maintenance in response to injury and aging. (A) *Left*, PLM neurons imaged 24 h following injury. All three groups are in the *zdIs5 (Pmec-4-GFP)* transgenic background. The red arrows indicate where the axon was severed. *Right*, axon regrowth measured 24 h following axotomy. The length of regrowing axons was normalized to the average of control. The numbers above the graph represent the number of PLMs tested in axon regrowth assay. non-parametric Mann-Whitney test for comparison between control and *ept-1*. Kruskal-Wallis test followed by Dunn’s post hoc test for comparison of *ept-1* and rescue groups. (B) *Left*, z-stacked confocal images of PLM axons in 5 day old adults. *Right*, quantification of PLMs with axon degeneration. The number of PLMs scored for each genotype is indicated above the corresponding bar. Fisher’s exact test. Scale bars = 10 µm. \**p*<0.05 and \*\**p*<0.01.

### Loss-of-function in *dip-2* bypasses requirements for *cept-2* or *ept-1* in axon regrowth

We previously found that loss-of-function mutation in *C. elegans dip-2* leads to progressive formation of ectopic axon neurites in multiple neuron types, as well as enhanced TRN axon regeneration following injury (Noblett *et al*. 2019). To test for genetic interaction between *dip-2* and the Kennedy pathway, we constructed *dip-2(0) cept-2(0)* double mutants and examined axon regrowth. Different allele combinations of double mutants exhibited similar or even more enhanced axon regrowth compared to *dip-2(0)* single mutants (Figure 5A,B). As *dip-2* and *ept-1* are ∼150 kb apart on chromosome I, we engineered *ept-1* deletions in the *dip-2(0)* background to generate *dip-2(0) ept-1(0)* double mutants (Figure 3B; Figure 5A,B). Such double mutants exhibited partial but significant suppression of *ept-1(0)* body length phenotypes (Figure S3A). However, the double mutants had brood sizes comparable to that in *ept-1* single mutants, indicating that loss-of-function in *dip-2* does not suppress *ept-1* requirement in fertility (Figure S3B).

**Figure 5.**
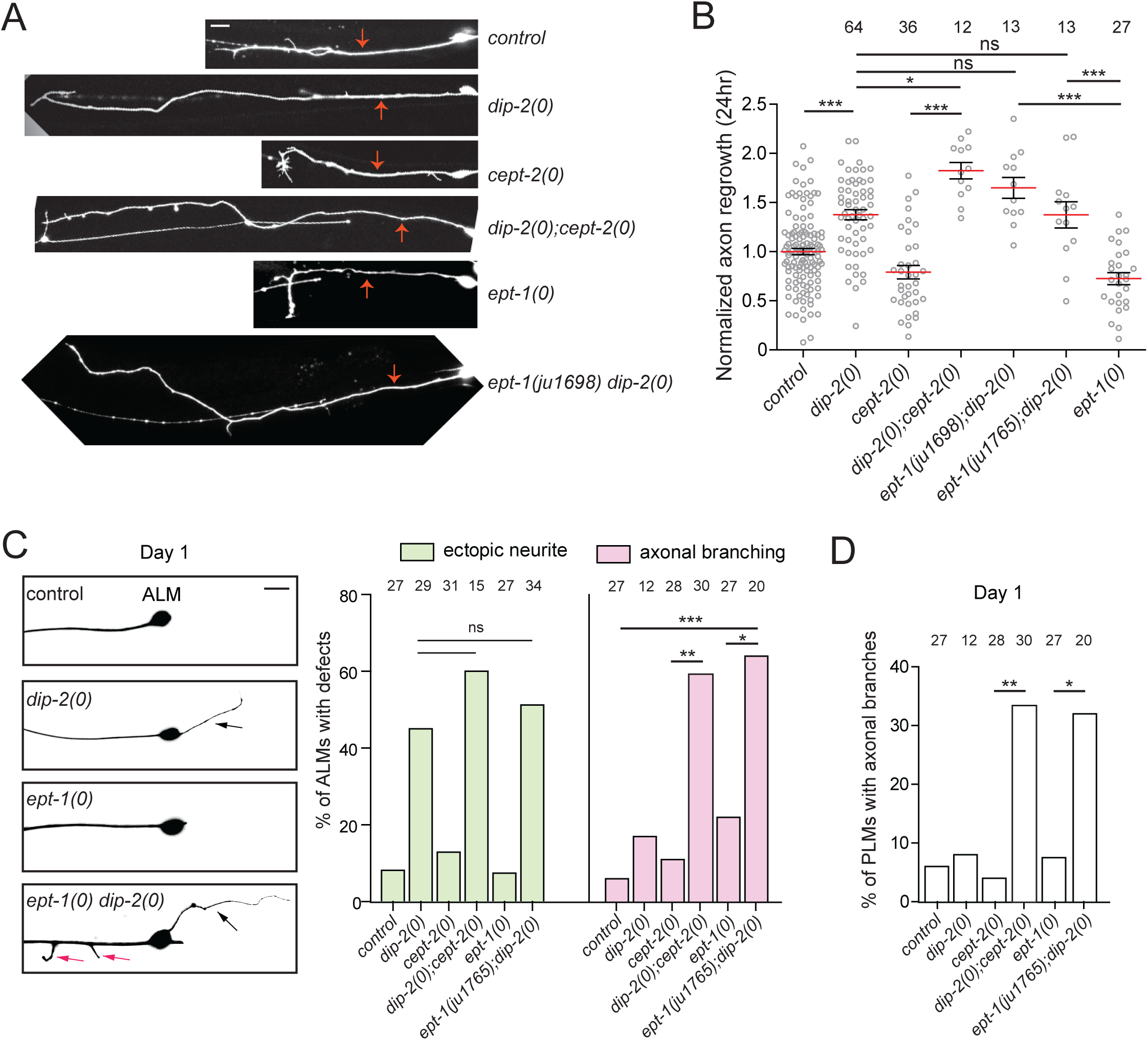
*dip-2* bypasses the requirement of *cept-2* and *ept-1* in axon regrowth. (A) PLM neurons imaged 24 h following injury. The red arrows indicate where the axon was severed. All six groups are in the *zdIs5 (Pmec-4-GFP)* transgenic background. (B) PLM axon regrowth measured 24 h following axotomy. The length of regrowing axons was normalized to the average of control. The numbers above the graph represent the number of PLMs tested in axon regrowth assay. (C) *Left*, Images of ALM at 1-day old adult stage. *Right*, Quantification of ectopic neurite outgrowth from the ALM soma and branching from the ALM axon. ALM posterior neurites >10 µm were scored as ‘‘ectopic’’. The number of ALMs scored for each genotype is shown above the corresponding bar. Fisher’s exact test. (D) Quantification of PLMs with axonal beadings in 1-day old adults. The number of PLMs scored for each genotype is indicated above the corresponding bar. Scale bars = 10 µm. ns means not significant. Fisher’s exact test. \**p*<0.05, \*\**p*<0.01, and \*\*\**p*<0.001.

Our results suggested that loss-of-function in *dip-2* may bypass the requirement for *cept-2* or *ept-1* in axon regrowth. We further asked whether *dip-2* interacts with other genes involved in lipid metabolism or signaling (Figure S3C,D). In mammalian optic nerve neurons, injury elevates levels of Lipin1 and DGAT, driving synthesis of triacylglycerol (TAG) at the expense of phospholipids and contributing to the low endogenous level of regeneration in such neurons; loss-of-function in Lipin1 or overexpression of Kennedy Pathway genes increases axon regeneration due to enhanced phospholipid synthesis (Yang *et al*. 2020a). DIP-2 proteins generate acyl-CoAs which could act as a source of fatty acids for Lipin1-catalyzed conversion of DAG to TAG, contributing to inhibition of axon regrowth. Loss of function in *lpin-1*, the *C. elegans* ortholog of Lipin1, causes early larval lethality (Golden *et al*. 2009). We therefore tested whether DIP-2 interacted with C-terminal domain (CTD) phosphatase CTDNEP1, which activates Lipin1 via dephosphorylation (Han *et al*. 2012). Disruption of *cnep-1* (pka *scpl-2*) the *C. elegans* ortholog of CTDNEP1, does not affect PC or PE levels, although PA and PI levels increase (Bahmanyar *et al*. 2014). *cnep-1(tm4369)* mutants exhibited normal TRN morphology and slightly reduced axon regrowth though it was not statistically significant (Figure S3D), suggesting *C. elegans* axon regeneration is less sensitive to loss of Lipin1 pathway function. Moreover, *dip-2(0) cnep-1(0)* double mutants resembled *dip-2* single mutants in axon morphology and TRN regrowth (Figure S3C,D). Overall, loss of function in *cnep-1* did not display clear genetic interactions with *dip-2.*Further, we tested the fatty acid amide hydrolase *faah-1*, which is involved in anandamide lipid signaling and promotes motor neuron regeneration (Pastuhov *et al*. 2012). Loss of function in *faah-1(tm5011)* did not significantly affect TRN morphology or regeneration and did not suppress *dip-2* phenotypes (Figure S3C,D). Taken together, these results suggest the TRN morphology and regeneration roles of *dip-2* may not involve altered anandamide pathways.

### Synergistic interaction between *dip-2* and *cept-2* or *ept-1* in axon branching

As DIP-2 also represses ectopic neurite outgrowth from the ALM soma, we also assessed neuronal morphology in the double mutants of *dip-2* with *cept-2* or *ept-1* at the day 1 adult stage and found that these mutants exhibited similar level of ectopic neurite outgrowth as *dip-2* single mutants (Figure 5C). However, the formation of axonal branches significantly increased in the double mutants of *dip-2* with *cept-2* or *ept-1* in both ALMs and PLMs (Figure 5C,D), suggesting synergistic interactions between the two parallel pathways in axonal branching. Collectively, our results suggest that *dip-2* and the Kenney pathway genes interact differently in axon regrowth and morphology maintenance and development/reproduction.

## Discussion

### The Kennedy pathway is required for axon regeneration in response to injury

Our results shed new light on the roles of specific phospholipid synthetic enzymes in axon maintenance and axon regeneration *in vivo*. Deletion of either *cept-2* or *ept-1* led to defects in axon regeneration (Figures 1D,4A). *cept-2* was especially required for axon extension rather than growth cone formation following injury and mildly enhanced axon regrowth when overexpressed (Figure 1D, S1C), suggesting phospholipid synthesis may be neuroprotective. The phenotypes of *cept-2* mutants may reflect depletion of both PC and PE as CEPT-2 is the *C. elegans* ortholog of CEPT1, which catalyzes the last step in both PC and PE branches of the Kennedy pathway. However, the phenotypes of *ept-1* mutants suggest PE biosynthesis could be specifically rate-limiting in axon regrowth in *C. elegans*. We find *ept-1* null mutants are subviable, possibly due to the ability of CEPT enzymes to catalyze some level of PE synthesis, or to PE synthesis from other cellular pathways such as PSD1. Deletion of *ept-1* may partly reduce PE levels below a threshold necessary for efficient membrane production in axon regrowth. In contrast, knocking down the Kennedy pathway genes in mice had no effect on injured axons both *in vitro* and *in vivo*, except in the Lipin1 mutant background (Yang *et al*. 2020a). The discrepancy could have arisen from the difference between global knockout in our study and cell/tissue-specific knockdown approaches taken in mice. It will be interesting to carry out future studies to determine whether disrupting phospholipid biosynthesis in a non-cell autonomous manner affects neurons upon injury. Nevertheless, our results reinforce the conclusion that augmenting Kennedy pathway function can improve axon regeneration. It will be important to understand how phospholipid biosynthesis can be precisely regulated to optimize repair while preventing unnecessary overgrowth.

### Phospholipid biosynthesis is neuroprotective in aged animals

Loss-of-function in either *ept-1* or *cept-2* resulted in normal TRN morphology in young adults but progressive axon defects in older animals, suggesting phospholipid levels may become critical in axon maintenance with age. Indeed, phospholipid levels decrease in human brains during normal aging (Hancock *et al*. 2022), and multiple studies have reported depletion of phospholipids in the brain of senior patients with Alzheimer’s disease (Pettegrew *et al*. 2001; Kao *et al*. 2020; Akyol *et al*. 2021). *cept-2* mutants specifically exhibited progressive and highly penetrant axonal swellings containing vesicular-like structures. We have not characterized these structures; however, they could correspond to membranous organelles or lipid droplets. Mutation of *atgl-1* leads to the formation of ring-shaped lipid droplets in most neurons including TRNs (Yang *et al*. 2020b) whereas the vesicular-like structures observed in our studies are slightly irregular in shape. As a mitochondrial marker displays discrete puncta along the axon in touch neurons (Knowlton *et al*. 2017), the vesicular-like structures could contain other membrane-bound organelles such as mitochondria and lysosomes. In addition to these axonal swellings, nicks or breaks in axons were also observed in older *cept-2* adults, suggesting phospholipid synthesis is important in maintaining axon integrity.

Our findings of axon degeneration in 5 day old *ept-1* mutant adults echo the light-dependent retinal degeneration phenotypes resulting from knockdown of *Ept* in *Drosophila* (Midorikawa et al. 2010). Our findings also parallel the results from a study in which loss of function mutation in *pect*, the gene encoding the ECT ortholog, resulted in adult-onset axon terminal degeneration in *Drosophila* (Tsai et al. 2019; Zhao and Wang 2020). ECT is the rate-limiting enzyme in the PE branch (Infante 1977). Lipidomic analysis indicated *pect* mutants lack a subset of PE with long chain fatty acid (Tsai *et al*. 2019). Possibly the levels of long chain FA PE may be critical for axon integrity. In *Drosophila*, axon degenerative defects appear to result not only from reduced PE levels but from consequent activation of the SREBP pathway and induction of SREBP target genes such as tetraspanins (Tsai *et al*. 2019). *C. elegans* contains two ECT orthologs, *pcyt-2.1* and *pcyt-2.2*, that do not show overt neuronal phenotypes as single mutants (Kim *et al*. 2018); we have not investigated possible redundancy of *pcyt-2.1* and *pcyt-2.2* function. Conversely, *Drosophila* encodes two EPT orthologs, the function of which is not yet investigated. In the future, it should be of interest to systematically test the requirement for phospholipid synthesis for maintenance of specific neuron morphologies.

The Kennedy pathway genes (EAK, PECT) have been shown to function cell autonomously in Drosophila dendrite morphogenesis (Meltzer *et al*. 2017). Phospholipid synthetic enzymes (PCYT1 and PCYT2) have been shown to function cell autonomously in murine retinal ganglion cell regeneration (Yang *et al*. 2020a). In *C. elegans*, transcriptomic studies indicate Kennedy pathway genes are widely expressed though mRNA levels of *cept-2* and *ept-1* are relatively low in touch neurons compared to BDU neurons which form gap junctions with PLMs (Taylor *et al*. 2021). TRN-specific *cept-2* expression led to restoration of axon integrity in *cept-2* aging neurons, consistent with prior studies that neurons have the cell autonomous capacity for *de novo* phospholipid synthesis for the maintenance of their morphology (Midorikawa *et al*. 2010; Tsai *et al*. 2019; Zhao and Wang 2020). Although our rescuing GFP::EPT-1 reporter was not detectable in the nervous system, EPT-1 may be expressed at very low levels in neurons. Alternatively, PE synthesis via EPT-1 may be required non-autonomously for axon maintenance.

### Genetic interactions between phospholipid synthesis and the lipid regulator DIP-2

Dip2 proteins contain Adenylate Forming Domains (AFDs) implicated in regulation of fatty acid metabolism (Nitta *et al*. 2017; Mondal *et al*. 2022). In *Drosophila* and mammals, lack of *dip2* does not have dramatic effects on phospholipid levels but elevates levels of diacylglycerol (DAG), the substrate for enzymes in the final step of PC and PE synthesis (Mondal *et al*. 2022). This surplus of DAG could be involved in the enhanced axon regrowth of *dip-2(0)* mutants. As the Kennedy pathway is regulated by highly redundant enzymes at each step, loss of function in any one Kennedy pathway enzyme may be suppressed by elevated DAG levels in *dip-2(0)* acting via other enzymes at the same step. Whether suppression by restoration of phospholipid levels, or by some other mechanism remains to be elucidated. While *dip-2* antagonistically interacts with *cept-2* or *ept-1* in axon regrowth, it exhibits a synergistic interaction with either gene in axon branching. These differential genetic interactions suggest the complex mechanism of lipid metabolism underlying axon regeneration and maintenance.

### Phosphatidylethanolamine levels and human neurological diseases

The *Drosophila pect* gene encoding the rate-limiting enzyme for the middle step of the Kennedy pathway is critical for PE homeostasis to prevent retinal degeneration (Tsai *et al*. 2019). Interestingly, only a subset of PE containing very long chain fatty acids (>C30) was significantly reduced in *pect* mutants, suggesting these PE species may be crucial for axon health. Our finding of axon regrowth defects in *ept-1* mutants, coupled with prior findings of axon degeneration in *Drosophila* ECT mutants, point to the essential nature of PE synthesis via the Kennedy pathway in neuronal maintenance and repair.

Mutations in Kennedy pathway genes have been implicated in numerous human genetic diseases, highlighting the critical nature of PC/PE synthesis (Tavasoli *et al*. 2020). Our observations also shed light on how PE dysregulation might cause neurological defects in humans. Notably, mutations in human EPT1 (Selenoprotein 1, SELENOI) cause hereditary spastic paraplegia (HSP) (Horibata *et al*. 2018; Li *et al*. 2023). One HSP pedigree contains a missense mutation in EPT1 that strongly reduced catalytic activity, although interestingly PE levels in blood were normal in these patients (Ahmed *et al*. 2017). While underscoring the redundant nature of PE biosynthesis pathways, these observations are consistent with mild reduction in PE or a subset of PE having severe consequences on neural function. A recent study has reported that stress-induced inactivation of PE biosynthesis can trigger cellular senescence, which is accompanied by morphological changes (Tighanimine *et al*. 2024). An increasing number of studies hint that neurons, despite being post-mitotic cells, can undergo senescence, and accumulation of senescent cells is observed in neurodegenerative diseases(Jurk *et al*. 2012; Baker and Petersen 2018; Herdy *et al*. 2022; Herdy *et al*. 2024). Better understanding of how PE biosynthesis is regulated under various conditions would contribute to developing potential therapeutic strategies against diseases.

---

**Figure S1.**
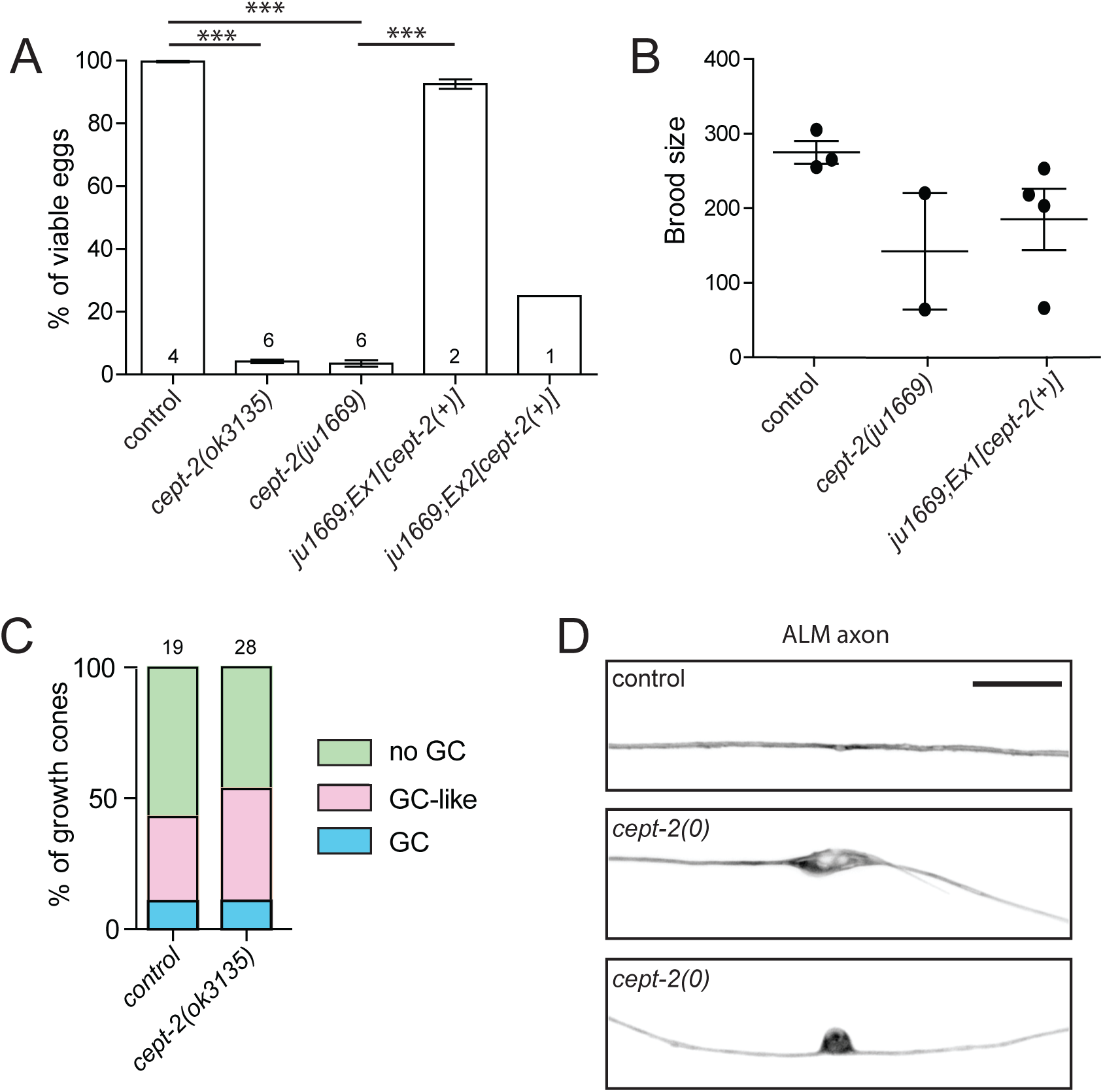
*cept-2* is required for development and reproduction. (A) Quantification of hatched animals from eggs laid by animals with various genotypes. The extrachromosomal arrays 1 and 2 (*juEx8005 and juEx8004*) harboring the *cept-2* minigene rescued embryo viability to different degrees. *juEx8005* was so stable that almost every animal contained the array. (B) Mean brood size measured over 5 days of adulthood. Extrachromosomal array 1 (*juEx8005*) rescued brood size. (C) Quantification of growth cone (GC) formation 24 h post axotomy. GC is defined as a structure with filopodia at the tip of a regenerating axon whereas GC-like is defined as a terminal swelling without filopodia. The numbers above the bars represent the number of PLM axons scored. (D) Confocal images of ALM axon imaged in 5-day old adults. Scale bar = 10 µm.

**Figure S2.**
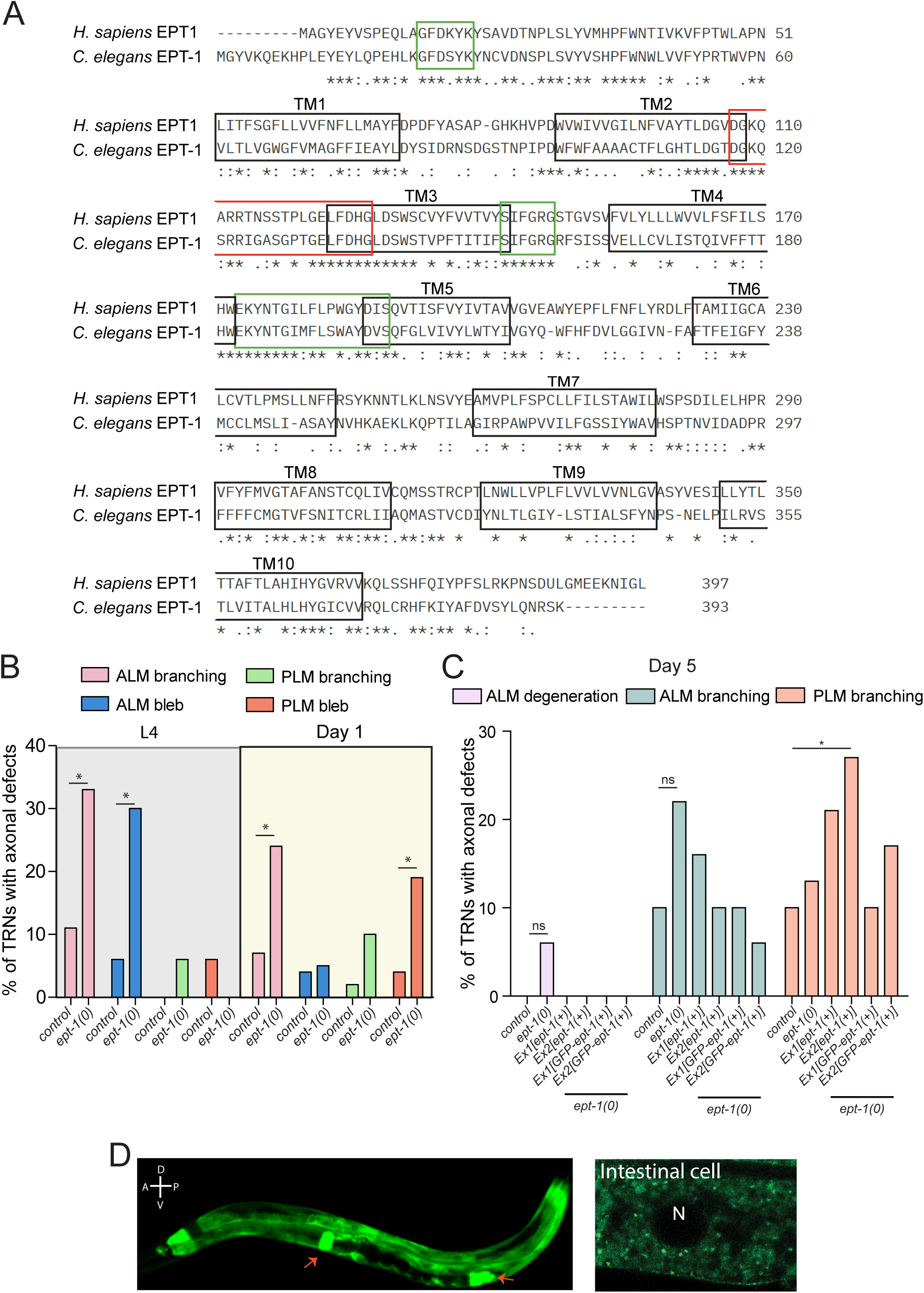
*ept-1* is required for reproduction and is expressed in the intestine and spermatheca. (A) Sequence alignment between human EPT1 and *C. elegans* EPT-1. Asterisk (*), colon(:), and period (.) represent residues with full conservation, partial conservation between amino acid groups of similar properties, and weak conservation between amino acid groups of weakly similar properties, respectively. Black-boxed regions indicate the 10 transmembrane regions (TMs), which were predicted using DeepTMHMM (Hallgren et al., 2022). Red- and green-boxed regions indicate the signature motif conserved in all CDP-alcohol phosphatidyltransferases and conserved residues specific to the EPT-1 orthologs, respectively. (B) Quantification of axonal defects in ALMs and PLMs at L4 and day 1 adult stages. N= 18-45. Fisher’s exact test. \**p*<0.05. (C) Quantification of various axonal defects in ALMs and PLMs at the day 5 adult stage. N=10-62. (D) z-stack of 15 confocal images of the entire worm (*left*) and z-stack of 3 confocal images of an intestinal cell (*right*) expressing GFP-tagged EPT-1 under the control of *ept-1* promoter at the day 1 adult stage. GFP::EPT-1 displayed punctate expression in the cytoplasm of intestinal cells. Red arrows point to spermatheca. N in white indicates nucleus.

**Figure S3.**
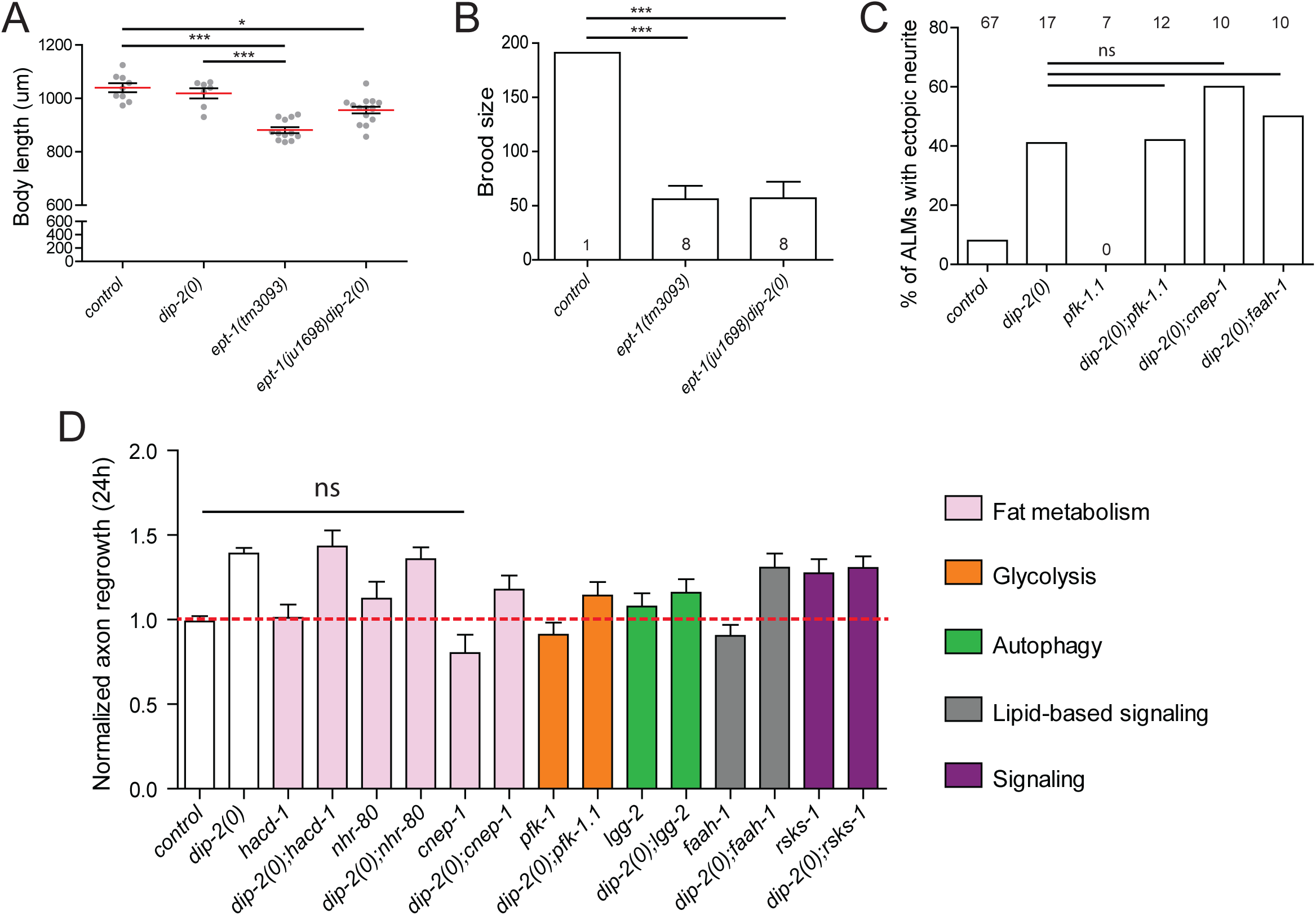
Genetic interaction between *dip-2* and lipid related pathways. (A) Body length measured in day 1 adults. Kruskal-Wallis test followed by Dunn’s Multiple Comparison Test. (B) Mean brood size measured for 5 days of adulthood. The numbers indicate the number of animals analyzed. One-way ANOVA and Tukey’s post hoc test. (C) Quantitation of ectopic neurite outgrowth from the ALM soma. The number of ALMs scored for each genotype is shown above the corresponding bar. Fisher’s exact test. (D) PLM axon regrowth measured 24 h following axotomy. The length of regrowing axons was normalized to the average of control. N=8-30. Fisher’s exact test. ns=non-significant. \**p*<0.05 and \*\*\**p*<0.001.

## Experimental Procedures

### Genetics and strain construction

Wild-type *C. elegans* used in this study is the N2 Bristol variant (Brenner 1974). Unless otherwise stated, strains were maintained at 20°C on Nematode Growth Media (NGM) plates seeded with *E. coli* OP50. Strains were constructed using standard procedures which include crossing using males, visually detecting behavioral and/or morphological phenotypes, and genotyping for mutant alleles. *ept-1(tm3093)* mutants were obtained from Mitani lab (Tokyo Women’s Medical University School of Medicine) as a balanced stock and later outcrossed to generate a homozygous strain.

### CRISPR gene editing

CRISPR editing used a *dpy-10* co-CRISPR strategy (Arribere *et al*. 2014). To generate a deletion allele of *cept-2*, a *cept-2* crRNA, *dpy-10* crRNA, tracrRNA, and Cas9 protein (QB3 MacroLab, UC Berkeley) were injected into N2. *ju1669* animals laid mostly dead eggs and few hatched, phenocopying *ok3135*. We obtained a homozygous strain and sequenced the region targeted by the *cept-2* crRNA; 4 nucleotides were removed from exon 2, resulting in a frame shift and a premature stop codon. To generate additional deletions of *ept-1* in the *dip-2(gk913988)* background, two *ept-1* crRNAs, *dpy-10* crRNA, tracrRNA, and Cas9 protein were injected into CZ26881 [*zdIs5;dip-2(gk913988)*]. Several F_1_ Dpy animals were pooled into one plate and their F_2_ progeny were genotyped for the presence of large deletions.

### Transgene construction and germline transformation

To generate a *cept-2* minigene genomic construct, ∼2.5 kb spanning the intergenic region between *cept-2* and the upstream gene F47B8.10 was amplified by PCR from genomic DNA and fused to *cept-2* cDNA amplified from N2 cDNA library, in pCR8. This resulting plasmid was then used as a template to amplify the intergenic region and the first exon of *cept-2* by PCR and fused to the ∼2 kb PCR-amplified genomic region spanning from exon 2 to the 3’ UTR in pCR8, to generate clone pCZ1075. To generate an *ept-1* genomic construct, the entire genomic region encompassing the ∼1.5 kb intergenic region between *ept-1* and the upstream gene *gnrr-1*, all the exons and introns, and ∼0.5 kb 3’ UTR was amplified by PCR and inserted into pCR8, to generate clone pCZ1074. To generate a GFP-*ept-1* genomic construct, GFP was amplified by PCR and inserted between the promoter and the first exon in pCZ1074, to generate clone pCZ1076. To generate a *cept-2* TRN-specific expression construct, *cept-2* cDNA was subcloned into pCZGY603. All transgenes constructed in our study were injected at 10 ng/µl of the DNA of interest with 2 ng/µl of P*myo-2*-RFP and 90 ng/µl of pBluescript as filler DNA.

*cept-2* minigene: *juEx8005* arrays displayed almost 100% transmission whereas *juEx8004* array displayed 40% transmission. We have not established whether *juEx8005* is highly stabilized or chromosomally integrated.

### Quantitation of brood size

One L4 animal was placed in a plate, allowed to lay eggs for a day, and transferred to a new plate every day for four consecutive days, and the number of eggs laid in each of five plates was counted and combined to obtain the total number of eggs laid by the animal. To quantify the percentage of viable eggs, an L4 animal was single-housed, allowed to lay eggs for 2 days and then transferred to new plates 2 days in a row. The number of eggs laid on the first plate was counted and two days later the number of hatched animals were counted to calculate the percentage of viable eggs. The same procedure was done with the second and third plates where the animal laid eggs.

### Characterization of neuronal morphology

A Zeiss Axio Imager M2 compound microscope was used to score neuronal morphology in animals anesthetized using 2 mM levamisole in M9 buffer. TRN labelled with Pmec-4-GFP(*zdIs5*) was scored for morphology. In scoring ALM ectopic neurites, any projections from the ALM soma other than the anterior axon were measured and scored as “ectopic posterior neurite” if longer than 10 µm. Axonal swellings of diameter >2µm were scored as beadings.

### Microscopy

Animals were immobilized in a drop of M9 buffer with 2 mM levamisole and mounted on a 4% agar pad. All confocal fluorescence images were acquired with a PlanApochromat 63x/NA1.4 oil immersion objective lens using Zeiss LSM800 (Axio Observer.Z1/7) confocal microscope. Maximum intensity projections were performed using Fiji ImageJ. Bright field images of gross morphology of animals with any genotype were taken either on a Zeiss Axio Imager M2 compound scope or Zeiss LSM800 confocal microscope at 10X magnification under identical settings.

### Confocal settings and Airyscan details

For Zeiss Airyscan imaging, animals were immobilized with 10 mM levamisole and mounted on a 4 agar or 10% agarose pad. Images were acquired using a Plan-Apochromat 63x/NA 1.40 oil immersion objective using Zeiss LSM900 microscope (Axio Observer.Z1/7) equipped with Airyscan 2. In Zen 3.4(Blue) software, SuperResolution mode with ‘Best Signal’ detection setup was selected. The laser for green fluorescence was set at 488 nm (470-525), 1.5-5% laser power and 800 V detector gain. The laser for red fluorescence was set at 561 nm (550-617), 12-15% power, and 800 V detector gain. Scans were in frame mode with unidirectional scan direction and scan speed of 7. For Airyscan SR imaging we used autofilter with Super Resolution values from 7.5-8.4.

### Axon regeneration assays

PLM axons were severed at a distance of 40-50 µm from the soma using femtosecond laser on a spinning-disk confocal microscope as previously described (Wu *et al*. 2007). Animals were recovered and then imaged 24 h later to quantify the regrowth of severed axon on Zeiss LSM710 or LSM800 confocal microscope. Measured regrowth lengths were normalized to that of the same day control (*zdIs5*).

## Data availability

Strains and plasmids are available upon request. The authors affirm that all data necessary for confirming the conclusions of the article are present within the article, figures, and tables.

## Acknowledgements

We thank the members of the Jin and Chisholm labs for valuable input throughout the work, especially Zilu Wu for help with laser axotomy setup. We thank Dongyeop Lee for providing us with *yhEx279*. Some strains were provided by the CGC, which is funded by NIH (P40 OD010440).

